# A modified RGB trichrome (HemRGB) for improving nuclear staining: application to colorectal cancer histopathology

**DOI:** 10.1101/2023.12.04.569959

**Authors:** Francisco Gaytán

## Abstract

The RGB trichrome staining method has been used to highlight two major components of the extracellular matrix, collagen and glycosaminoglycans. While the RGB trichrome efficiently stains extracellular matrix components, it lacks a nuclear stain, limiting its application in histopathology. To address this issue, a modification of the original stain, named HemRGB trichrome, has been developed. This modification incorporates iron hematoxylin for improving nuclear staining while retaining specificity for the staining of extracellular matrix. The application of HemRGB trichrome staining to samples from both normal colonic tissues and colorectal adenocarcinomas (CRC) provides a robust nuclear staining, together with a high-contrasted staining of tumor microenvironmental components, such as infiltrating immune cells, collagen and ground substance, extracellular mucins, as well as contrasted interfaces between CRC metastases and liver parenchyma. This study underscores the potential of HemRGB trichrome as a valuable tool for histopathological studies, especially for cancer evaluation, where nuclear characteristics are particularly relevant.

## Introduction

Histopathological examination of tissue sections under the microscope continues to be the gold standard method to assess tissue alterations in various diseases. For this, adequate tissue staining is crucial for accurate tissue architecture analysis, and several stains are employed to visualize the different tissue constituents^**1**^. Hematoxylin and eosin (HE) staining, widely used as a routine histological method, provides a general overview of tissues, that could be complemented with additional stains for specific purposes. So, trichrome stains differentiate collagen from smooth muscle and are particularly useful in assessing collagen abundance^**1**^. The RGB trichrome was developed for the staining of two major components of the extracellular matrix^**2**^. Indeed, the RGB trichrome utilizes picrosirius red to stain collagen red and alcian blue to stain glycosaminoglycans blue, resulting in two non-overlapping colors. The third component of the RGB trichrome, fast green, acts as a general protein stain, enhancing the contrast of red stained collagen fibers, and staining the erythrocytes in intense green. However, a weakness of the RGB trichrome method is the lack of a nuclear stain. Nuclear morphology provides valuable information, and changes in nuclear size, shape, chromatin pattern, and staining properties are hallmarks of cancer and are essential for its histopathological evaluation^**3**,**4**^.

While cell nuclei are generally not stained with RGB trichrome (Fig.1A), cells in some tissues such as the epithelium of the seminiferous tubules (Fig. 1B), and some neurons in the retina (Fig. 1C) and peripheral autonomous nervous system (Fig. 1D) may exhibit partial nuclear staining. However, fine details and the chromatin patterns are not well-appreciated. This limitation could be partially circumvented by staining parallel sections with HE or RGB trichrome (Fig 2A,B). Even, if archival histological sections are already stained with HE and unstained sections are not further available, the HE-stained sections can be de-stained (i.e., after digitation) and re-stained with RGB trichrome, providing results equivalent to those in *de novo* stained sections (Fig. 2C,D). The best solution, however, is the simultaneous observation of cell nuclei and extracellular matrix characteristics in the same tissue section. To address this limitation, the present study introduces a modification of the RGB trichrome, that incorporates a nuclear dye to enhance nuclear staining while maintaining the specific staining of the extracellular matrix components. This allows a better observation of cell nuclei (Fig. 3) even in those tissues in which nuclei are partially stained with RGB trichrome.

**Figure 1.**
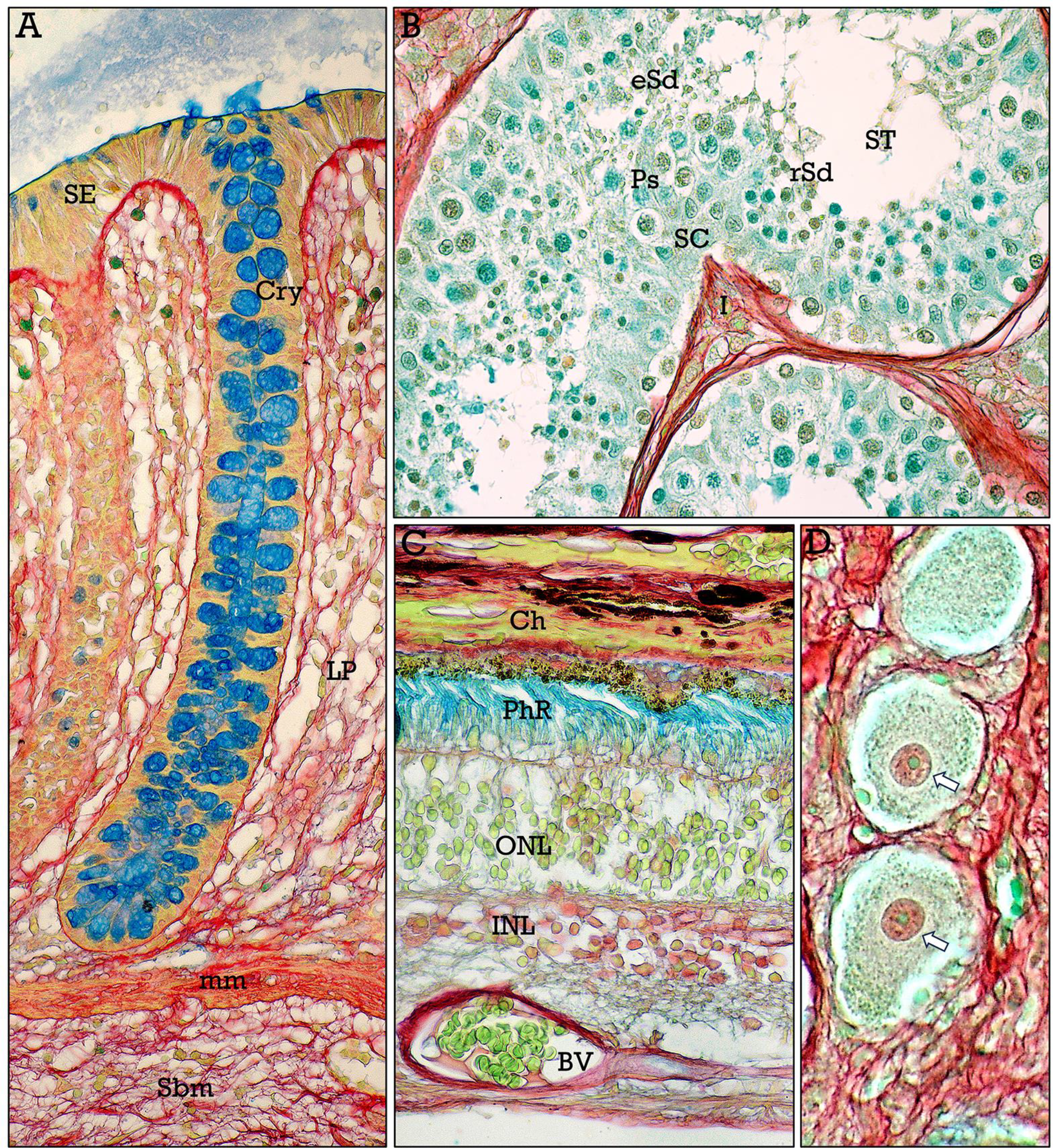
RGB trichrome-stained tissues. **A**, normal colonic mucosa showing the superficial epithelium (***SE***), a crypt (***Cry***) with abundant mucin-secreting (globet) cells, the lamina propria (***LP***), and the muscularis mucosae (***mm***) separating mucosa and submucosa (***Sm***). **B**, In the seminiferous tubules (***ST***) the nuclei of Sertoli cells (***SC***), pachytene spermatocytes (***PS***), and round (***rSd***) and elongated (***eSd***) spermatids can be identified. **C**, human retina showing the photoreceptor (***PhR***), the outer (***ONL***) and the inner (***INL***) nuclear layers, and retinal blood vessels (***BV***). The choroid (***Ch***) can be also observed. **D**, neurons of the autonomic nervous system with central nuclei and prominent nucleoli (***arrows***).

**Figure 2.**
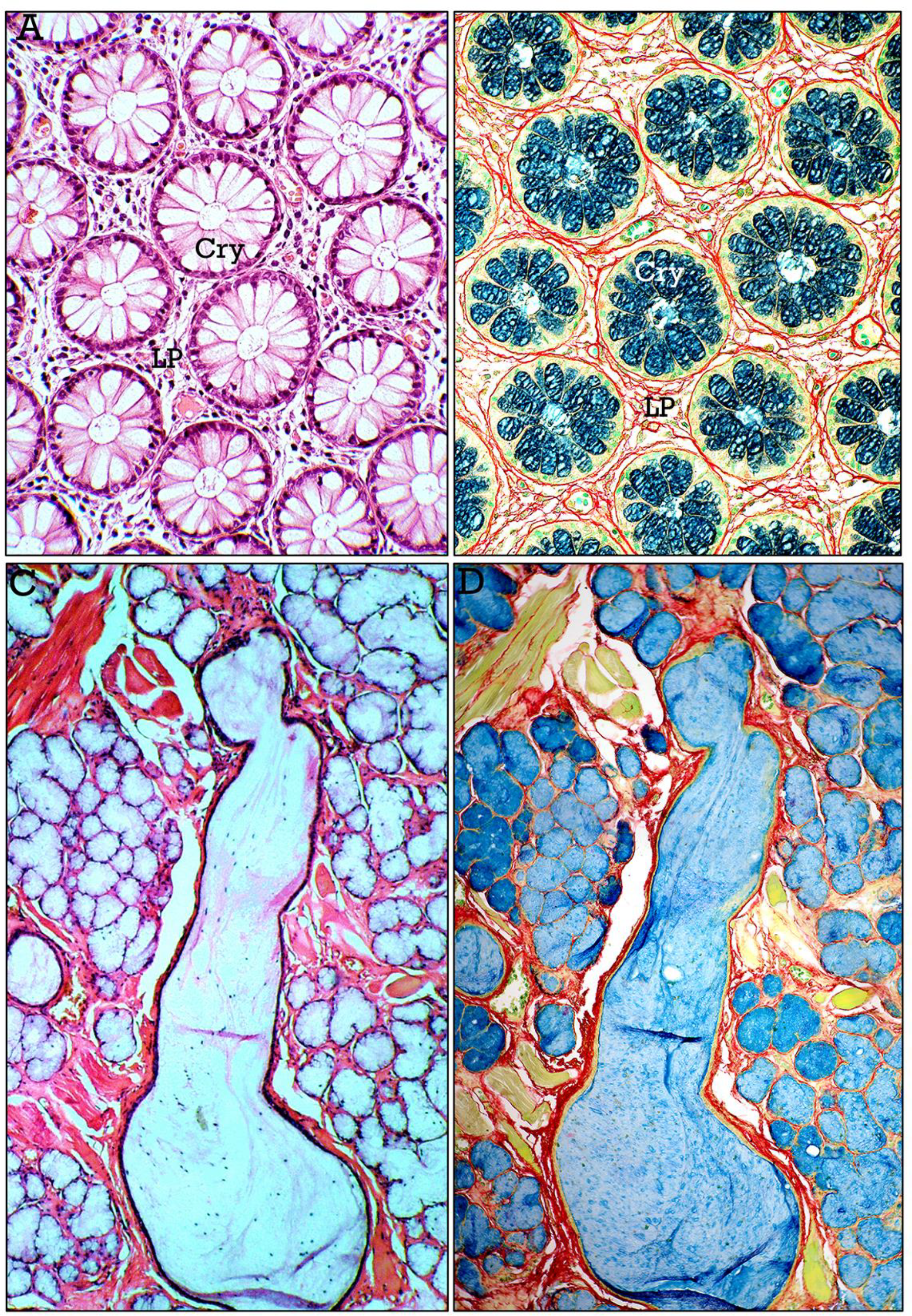
HE vs. RGB trichrome stains. **A**,**B**, parallel cross-sections of colonic mucosa stained with HE (**A**) or RGB trichrome (**B**), showing crypts (***Cry***) and the lamina propria (***LP***). **C**,**D**, The HE-stained section of salivary gland containing a pareidolic “lady in the garden” (**C**), was de-stained and re-stained with RGB trichrome (**D**).

**Figure 3.**
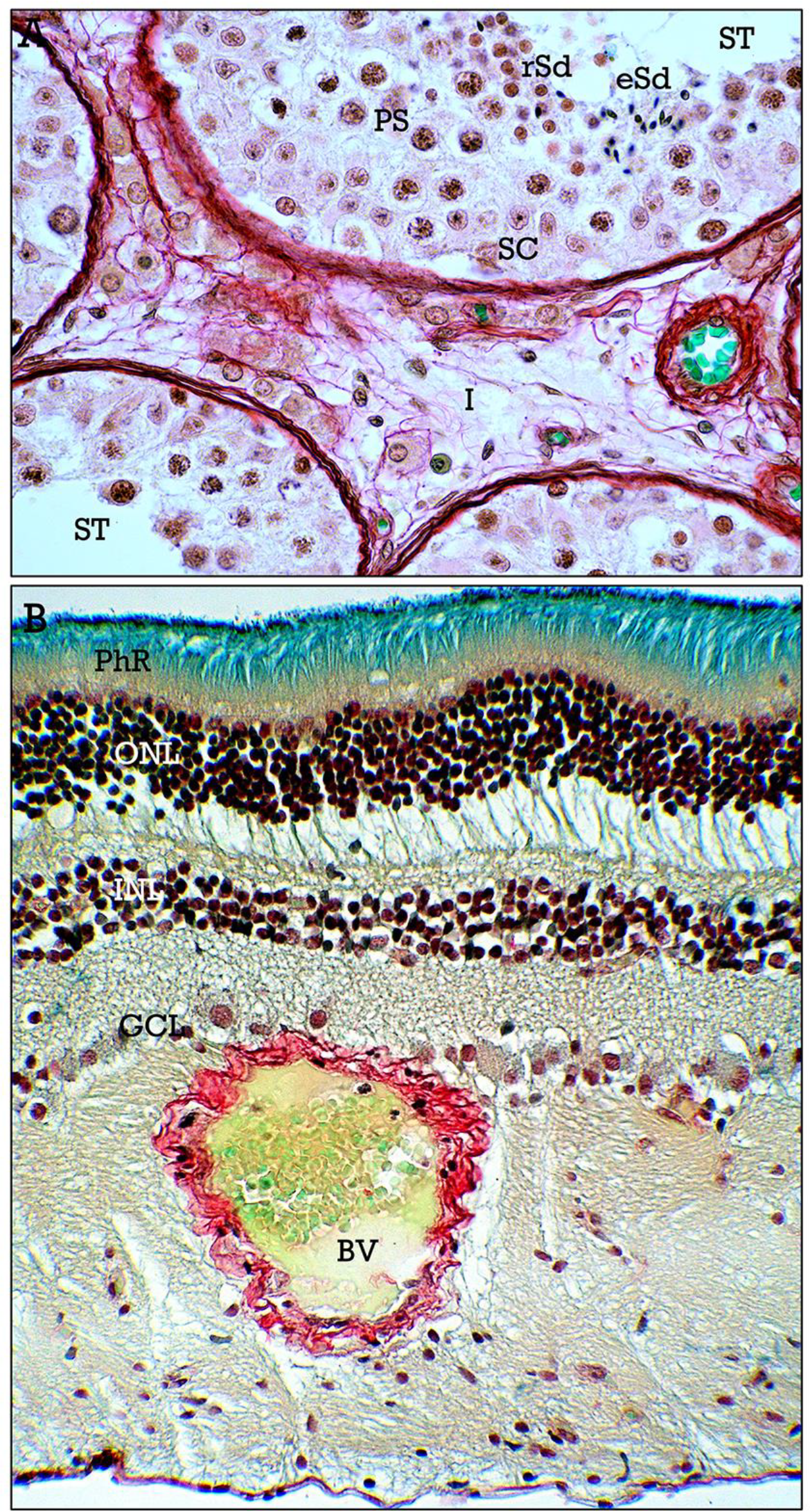
Enhanced nuclear visibility after HemRGB staining. **A**, seminiferous tubules (***ST***), showing enhanced staining of the nuclei of Sertoli cells (***SC***), pachytene spermatocytes (***PS***), round (***rSd***) and elongated (***eSd***) spermatids. Blood vessels and Leydig cells can be observed in the interstitial tissue (***I***). **B**, in the retina, the nuclei of neurons in the outer (***ONL***), inner (***INL***) and ganglionic (***GNL***) nuclear layers are intensely stained. Photoreceptor (***PhR***) layer and retinal blood vessels (***BV***) are clearly identified.

In this context, the aim of this study is to assess whether the modified trichrome stain, named HemRGB trichrome (acronym for Hematoxylin and RGB), is a reliable staining method for histological studies, particularly in the evaluation of cancer histopathology where nuclear characteristics are crucial^**5**,**6**^.

## Materials and Methods

Human tissue samples were obtained from the Bio-archive of the Department of Pathology of the Faculty of Medicine and Nursing. These specimens corresponded to tissues that were collected for diagnostic purposes between 1980-2000, adhering to contemporary legislation, including informed consent. The use of these samples was approved by responsible members of the Department of Pathology, granting patient confidentiality, and following strict adherence to current legislation. Tissues were archived as paraffin blocks.

Sections (5-6 μm thick) were cut, deparaffinized, hydrated through decreasing ethanol concentrations and submitted to HemRGB staining, which involved an initial staining with Weigert iron hematoxylin (Sigma-Aldrich, ref. HT1079-1SET) for 10 min following the provider instructions. After washing in tap water for 10 min, the sections were stained with RGB trichrome as previously described^**2**^. Briefly, the sections were sequentially stained with 1% (w/v) alcian blue in 3% aqueous acetic acid solution (pH 2.5) for 20 min, washed in tap water, stained with 0.04% (w/v) fast green in distilled water for 20 min, washed in tap water, and finally stained with 0.1% (w/v) sirius red in saturated aqueous solution of picric acid for 30 min, washed in two changes of acidified (1% acetic acid) tap water, dehydrated in 100% ethanol, cleared in xylene, and mounted with synthetic resin.

## Results

The staining properties of the HemRGB trichrome were tested on samples from colorectal adenocarcinomas. While detailed characterization of the different histotypes of colorectal cancer (CRC) was far beyond the study scope, colorectal adenocarcinomas were chosen to emphasize the importance of nuclear morphology, together with some other outstanding histological features relevant in histopathological assessment of CRC. Samples from the marginal areas of the resection pieces that were free of tumor and showed normal histology were used to assess staining characteristics of normal colonic tissues.

### Normal colonic tissues

Staining with HemRGB trichrome provided regular nuclear staining and highly contrasted tissue interfaces. The colonic mucosa (Fig. 4) showed a plain surface epithelium with absorptive and mucin-secreting (globet) cells, parallel cylindrical crypts containing abundant globet cells (Fig. 4A,B), and occasional enteroendocrine cells at the gland bottom (Fig. 4C). The connective tissue of the lamina propria contained abundant leukocytes. The mucosa was delimited by the muscularis mucosa. The submucosa (Fig. 5) contained connective tissue with abundant collagen, the nerves and ganglion cells of the submucosal plexus of Meissner (Fig. 5B), blood and lymphatic vessels and nerves (Fig. 5E). Lymphatic nodules, frequently observed in the mucosa, communicate with the submucosa through gaps in the muscularis mucosa (Fig. 5F). The most external layers of the colon corresponded to a continuous inner circular layer and a discontinuous external longitudinal layer of smooth muscle, containing the nerves and ganglion cells of the myenteric plexus of Auerbach (Fig. 5C).

**Figure 4.**
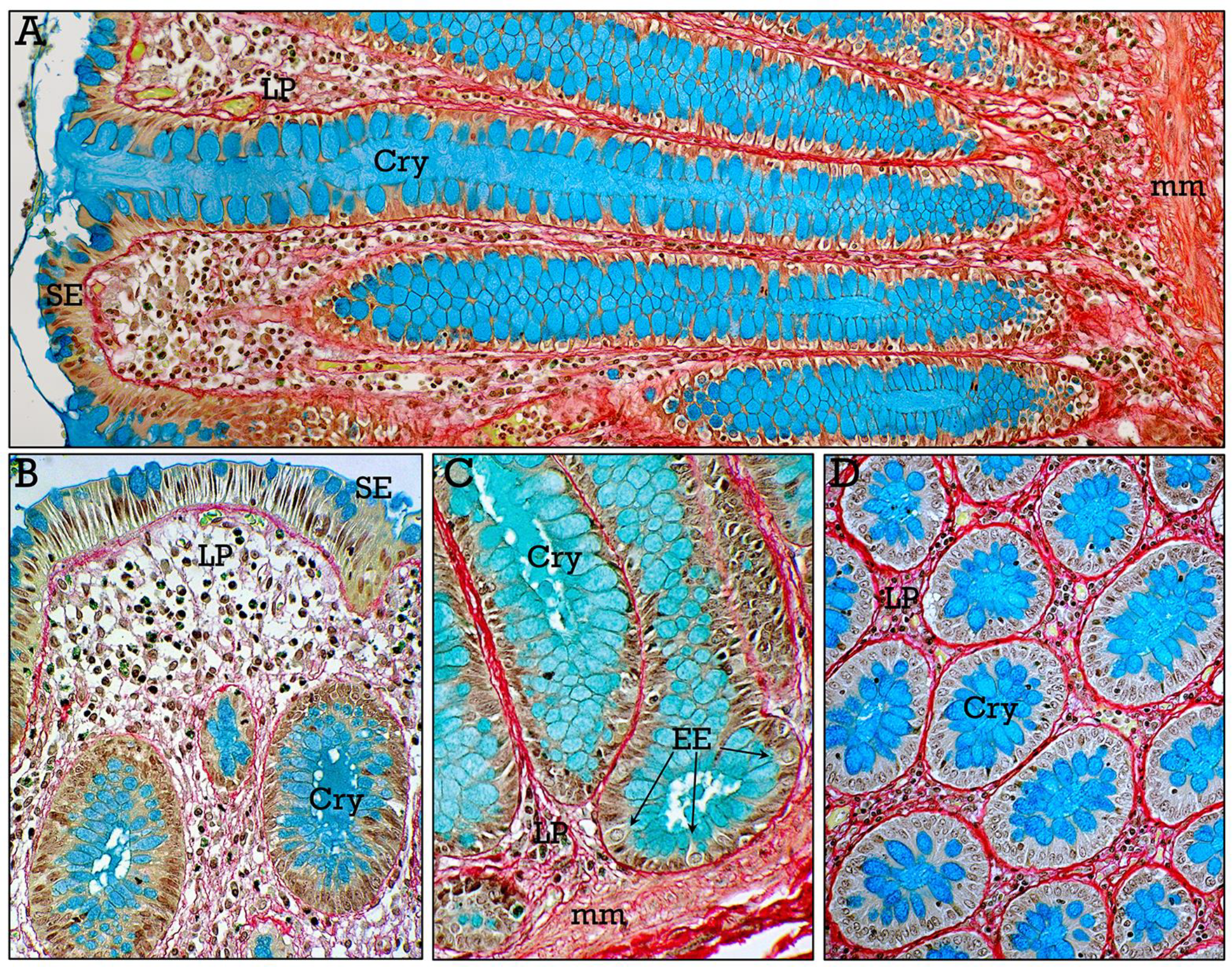
HemRGB-stained normal colonic mucosa. **A-C**, longitudinal sections of parallel tubular crypts (***Cry***) with prominent globet cells, extending from the surface to the muscularis mucosae (***mm***). The superficial colonic epithelium (***SE***), occasional enteroendocrine cells (***EE***) at the gland bottom, and the lamina propria (***LP***) with abundant immune cells, are shown at higher magnification in **B**,**C**. In **D**, transverse section of the colonic mucosa.

**Figure 5.**
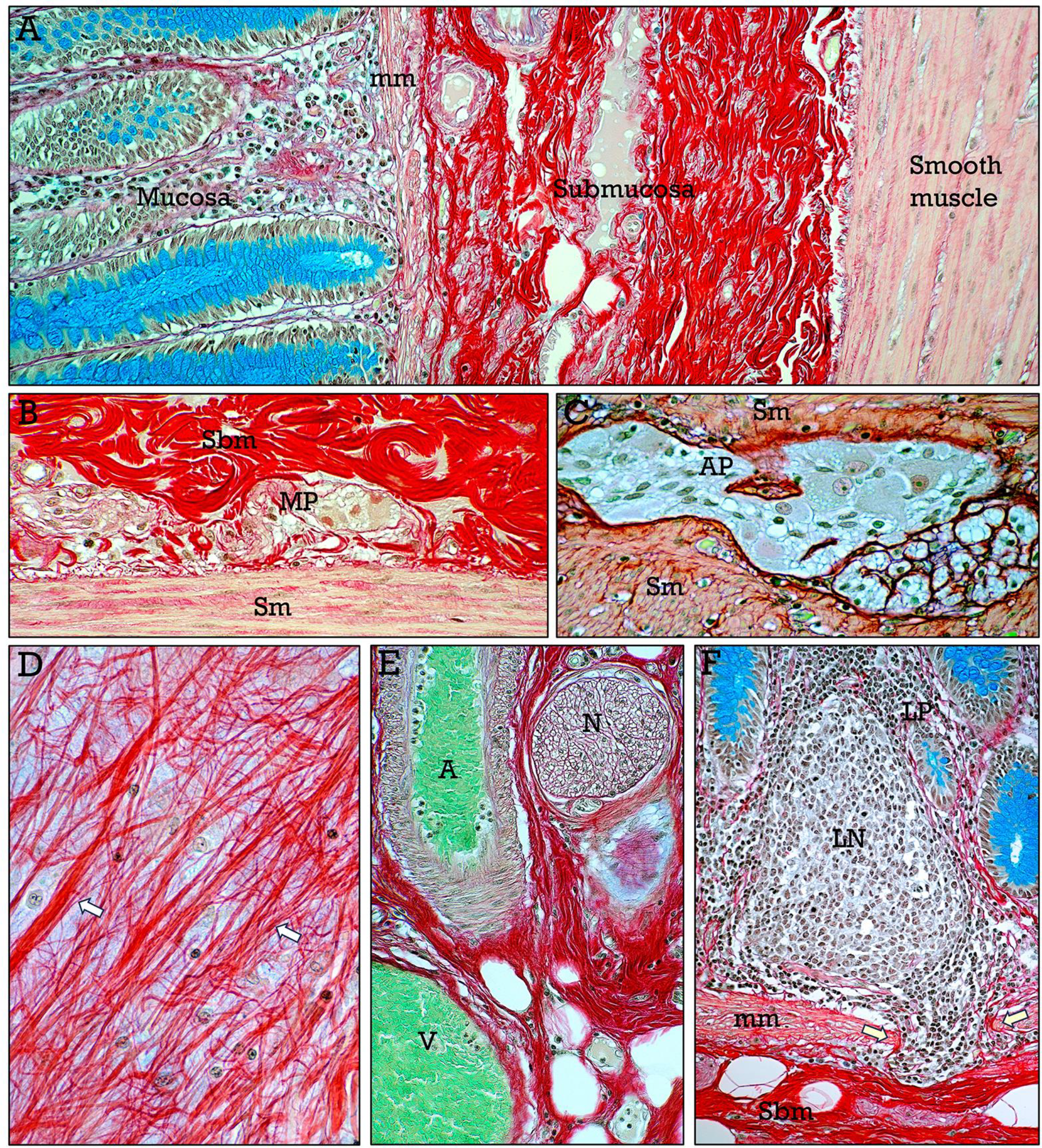
HemRGB-stained colonic wall. **A**, general view of the mucosa, submucosa and smooth muscle layers. Submucosal (**B**) and muscular (**C**) nervous plexuses of Meissner (***MP***) and Auerbach (***AP***) are shown. In the submucosa, collagen fibers are embedded in glycosaminoglycan background (**D**), and neurovascular bundles (**E**) are composed of branches of arteries (***A***), veins (***V***) and nerves (***N***). The lymphatic nodes (***LN***) in the mucosa (**F**) extend to the submucosa through gaps (***arrows***) in the muscularis mucosae (***mm***).

### Colorectal adenocarcinoma

Representative HemRGB tricrome-stained tissue sections from colorectal adenocarcinomas are shown in Figs. 6-11. Compared with normal mucosa, the adenocarcinomatous glands showed disorganized epithelium, decreased number of globet cells and abundant necrotic cell debris in the gland lumen (Fig. 6). All studied cases corresponded to invasive tumors (Fig. 7) infiltrating the submucosa (Fig. 7A) and muscular (Fig. 7B) layers. Morphological characteristics of CRC varied between zones, and different parameters such as nuclear morphology, immune cell infiltration, mucin secretion, and the amount of collagen and glycosaminoglycans in the tumor extracellular matrix, together with some liver metastases, were studied.

**Figure 6.**
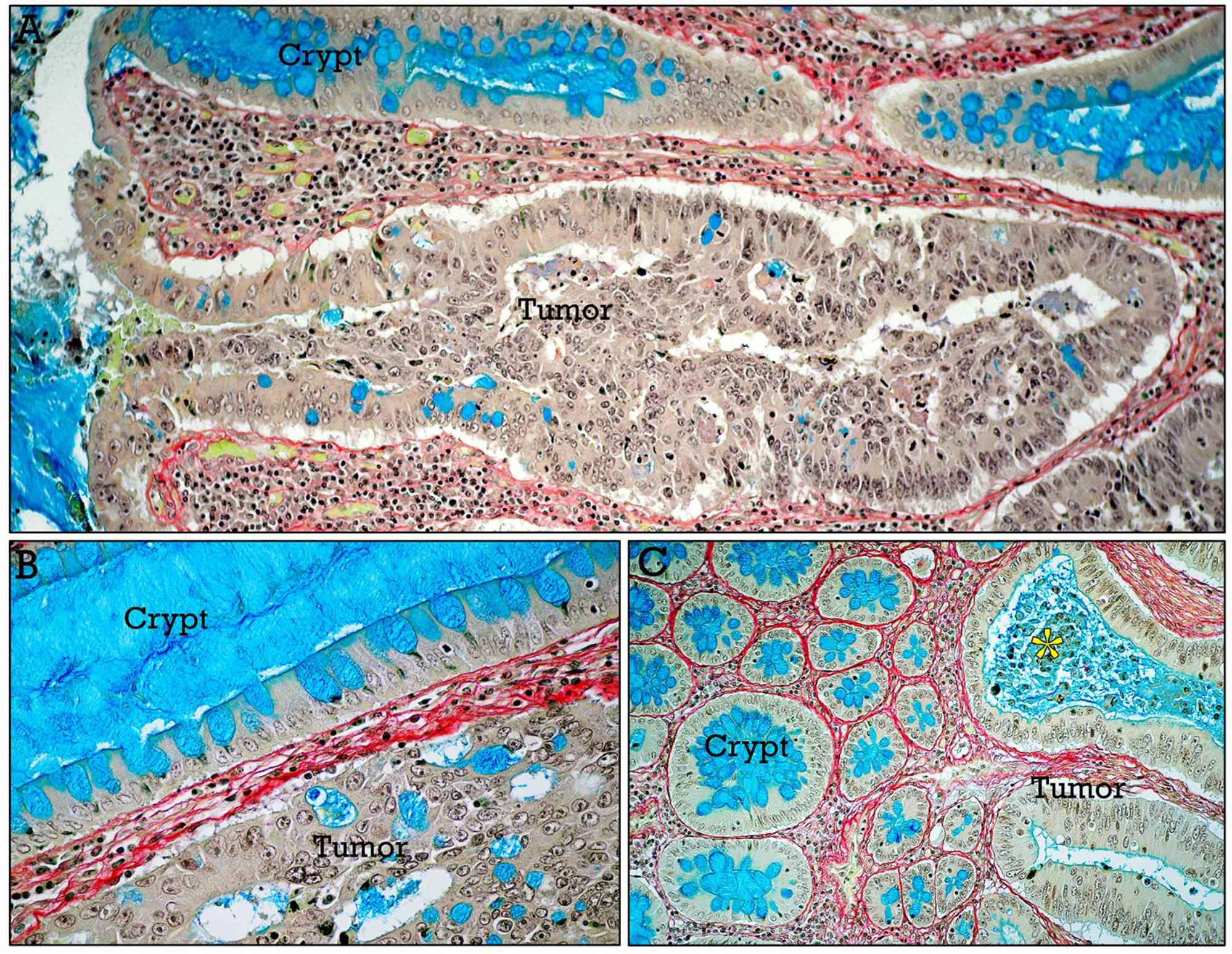
HemRGB-stained CRC. Longitudinal (**A**,**B**) and transverse (**C**) seccions of the mucosa containing adenocarcinomatous glands, showing disorganized epithelium, scarce mucin-secreting cells and abundant necrotic cell debris (***asterisk***), compared with normal crypts.

**Figure 7.**
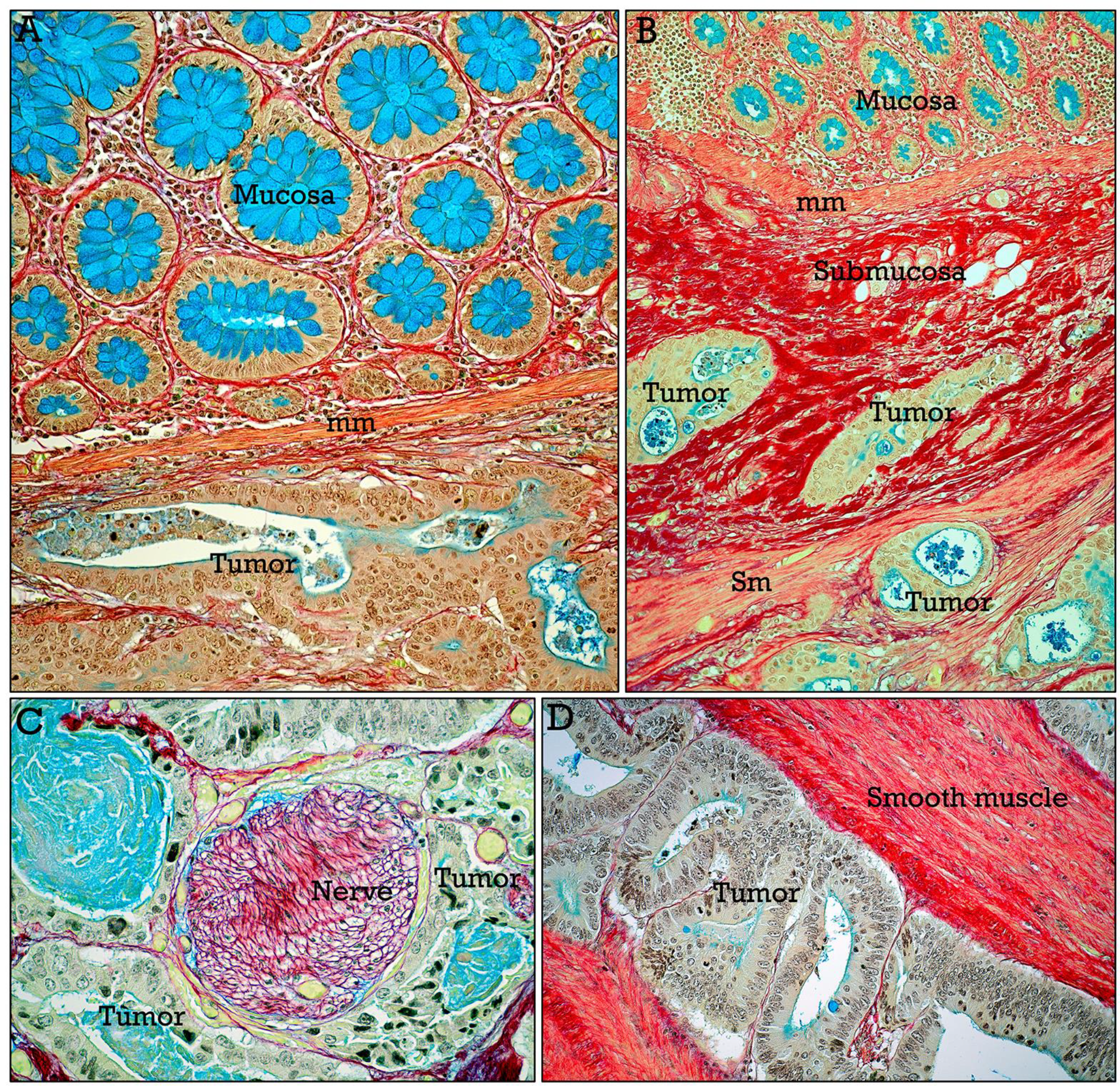
HemRGB-stained invasive CRC. Representative adenocarcinomas invading the submucosa (**A**) and the muscular layer (**B**). Perineural (**C**) and muscular (**D**) infiltration at higher magnification.

1. *Nuclear staining*. Representative nuclear staining in a sample from a poorly differentiated CRC is shown in Figure 8, exhibiting high nuclear pleomorphism, including atypical mitoses (Fig. 8B), apoptotic nuclei (Fig. 8C), prominent nucleoli (Fig. 8D), enlarged nuclei (Fig. 8E), and irregular nuclear shapes (Fig. 8F). The chromatin pattern was also evident.
2. *Immune cell infiltration*. The infiltration of the tumor by immune cells, primarily lymphocytes identified by their nuclear characteristics, varied across cases and was very abundant in some tumors (Fig. 9).
3. *Mucin secretion*. In general, mucin secretion was decreased in CRC, as evidenced by the scarcity of mucin-secreting cells even in the glands of well-differentiated tumors (Fig. 10A). However, in some areas, mucin-secreting cells displaying differential staining (blue or magenta) were occasionally observed (Fig. 10B). This resembles the reverse staining pattern of mucins in gastric intestinal metaplasia, where neutral mucins in the gastric mucosa stain red, while intestinal acidic mucins stain blue (Fig. 10C). Otherwise, mucins were very abundant in areas of tubulovillous glands (Fig. 10D), whereas the amounts of extracellular mucins range from small areas (Fig. 10E) to large mucin pools (Fig. 10F) in tumors with a prevalent mucinous component.
4. *Collagen and glycosaminoglycans in the tumor extracellular matrix*. Large differences existed in the amount of collagen in different tumors. In some cases, local areas of coarse collagen fibers were present (Fig. 11A), while a dense desmoplastic extracellular matrix, consisting of uniformly dense collagenous tissue, was present in some tumors (Fig. 11B). Glycosaminoglycans and proteoglycans were also occasionally abundant in the connective tissue and smooth muscle adjacent to the tumor (Fig. 11C,D). As a positive control for glycosaminoglycan staining, HemRGB staining of embryonic tissues was used (Fig. 11E).
5. *Liver metastases* Samples from livers containing CRC metastases were studied to assess the differential staining of metastatic and liver tissues, together with the characteristics of the tumor-liver interfaces. In normal liver tissue, the epithelium of biliary ducts showed apical mucin expression (Fig. 12A), which should not be confused with mucin-expressing CRC epithelial cells. Liver parenchyma (Fig. 12B) shows interconnecting plates of hepatocytes surrounding sinusoidal channels, and fine, red-stained collagen fibers are present in the space of Disse.

**Figure 8.**
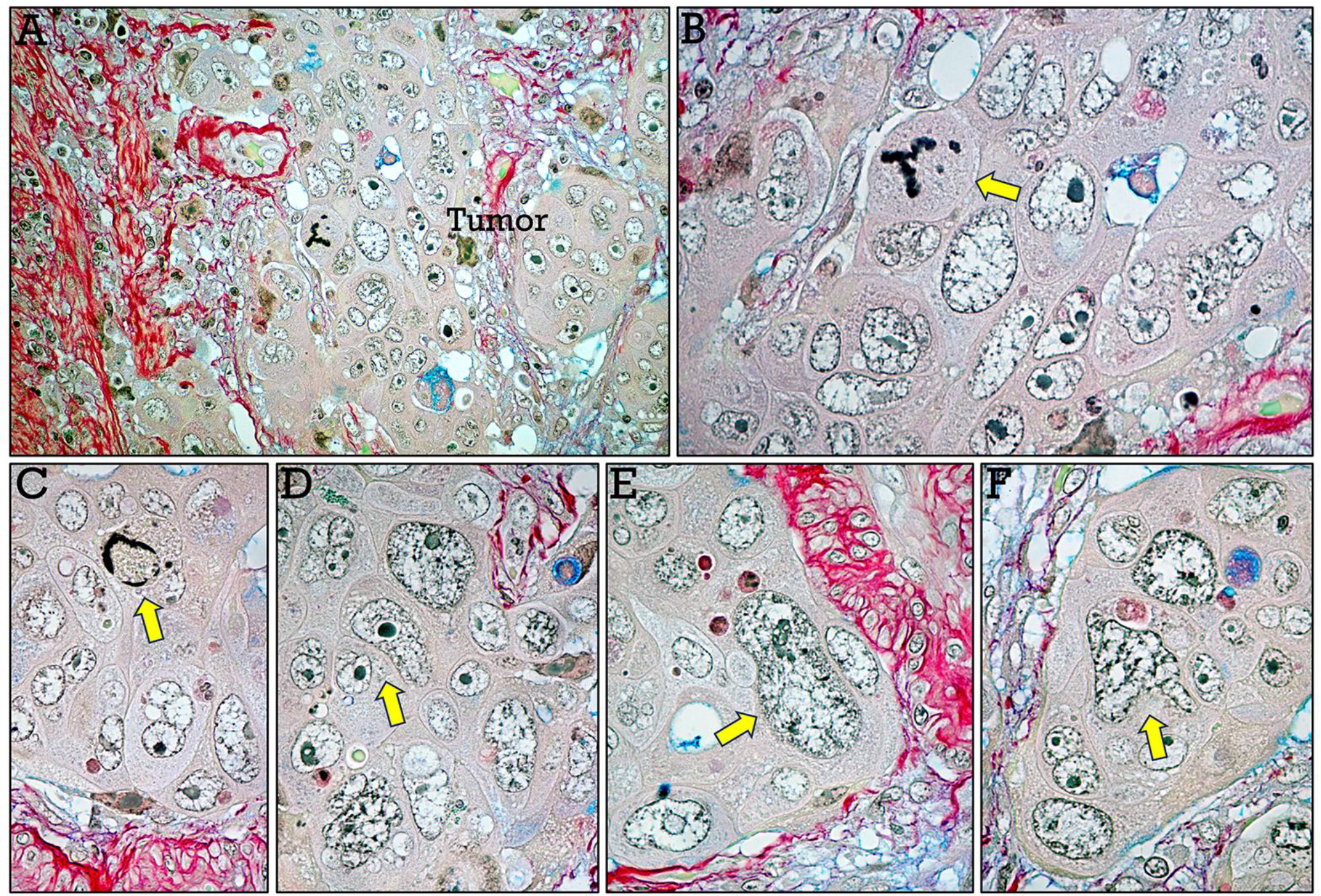
HemRGB-stained cell nuclei. Nuclear pleomorphism in a poorly differentiated CRC (**A**). Arrows indicate atypical tripolar mitosis (**B**), apoptotic nuclei (**C**), prominent nucleoli (**D**), enlarged (**E**) and irregularly shaped (**F**) nuclei.

**Figure 9.**
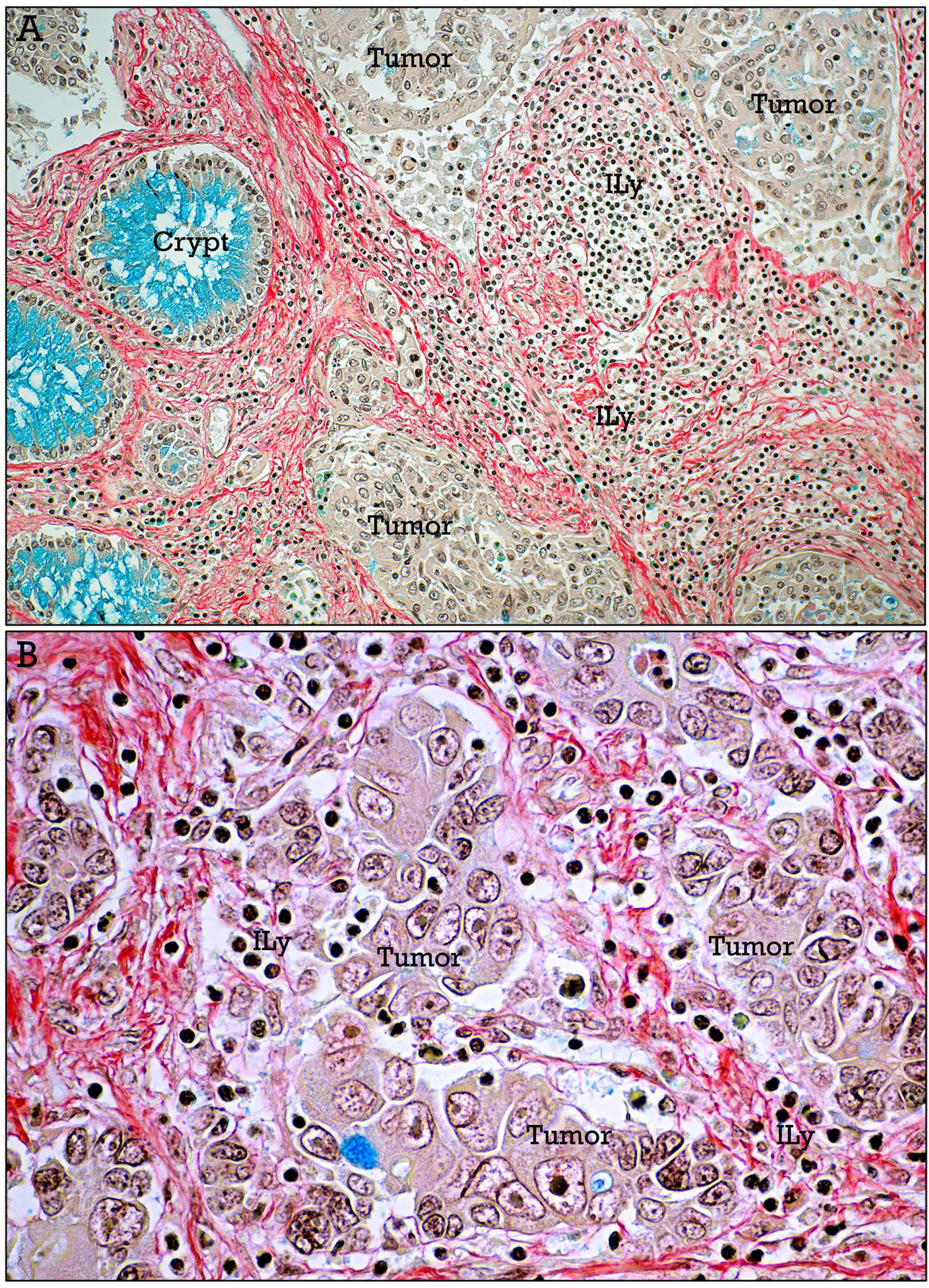
HemRGB-stained infiltrating leukocytes. **A**, abundant infiltrating lymphocytes (***ILy***), shown at higher magnification in **B**.

**Figure 10.**
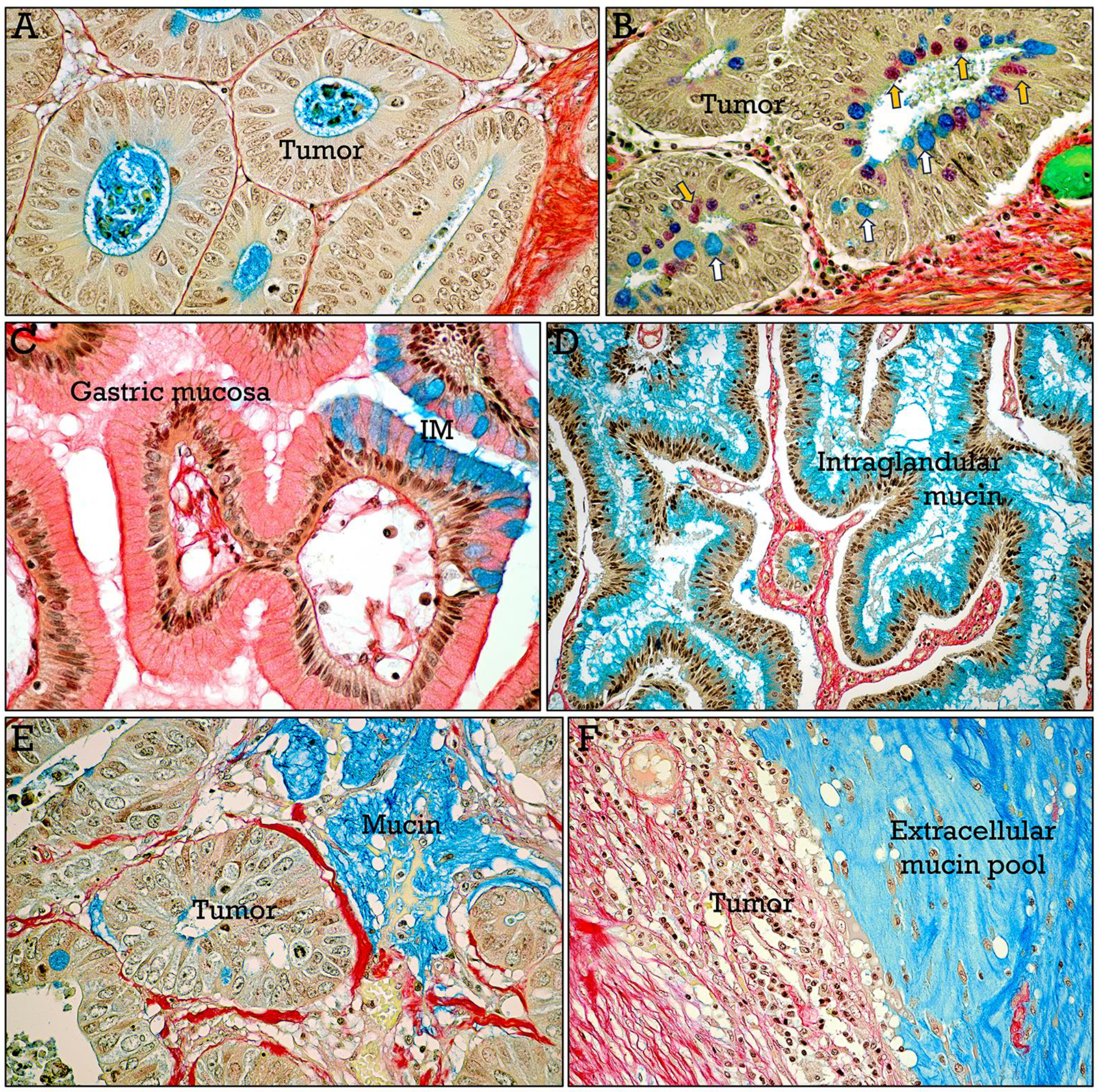
HemRGB-stained mucins. In general, decreased number of mucin-secreting cells were present in adenocarcinomatous glands (**A**). In some cases (**B**), differences in the staining properties of mucins were present, ranging from blue (***white arrows***) to magenta (***orange arrows***). This resembles the gastric mucosa (**C**) where neutral mucins are red stained, whereas acidic mucins in areas of intestinal metaplasia (***IM***) are blue stained. In **D**, abundant mucin in tubulovillous glands. Areas of extracellular mucins range from small (**E**) to large pools (**F**).

**Figure 11.**
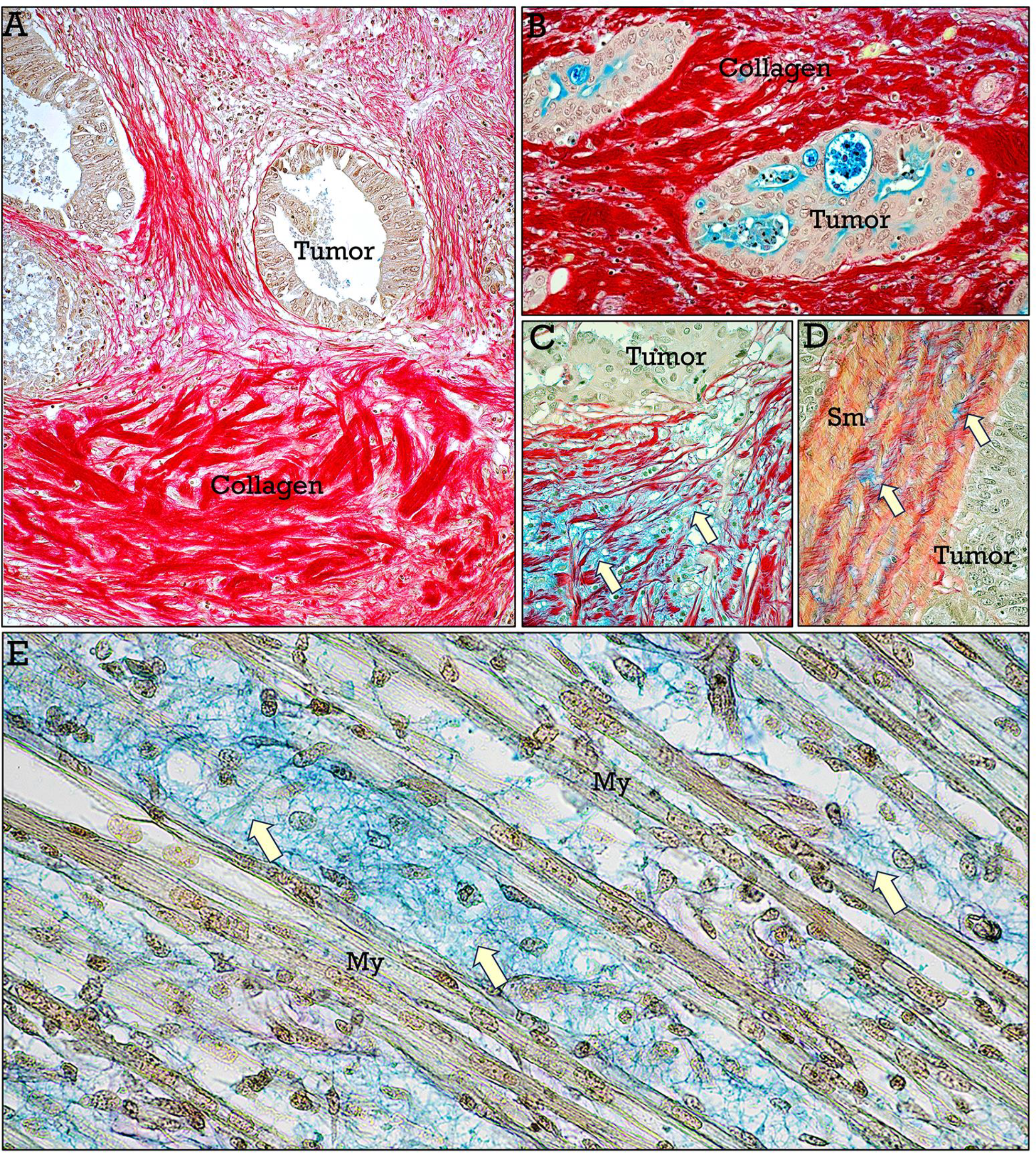
HemRGB staining of the tumor extracellular matrix. In some CRC, coarse bundles of intensely red stained collagen were present (**A**,**B**). Blue-stained glycosaminoglycans and proteoglycans were also abundant in the connective tissue matrix (**C**) and smooth muscle (**D**) in some CRC. Embryonic muscular tissue (**E**) showing myotubes (***My***) embedded in a glycosaminoglycan-rich, blue-stained matrix (***arrows***) used as a positive control.

Liver metastases were generally clearly differentiated from liver tissue. This was facilitated in some cases by the presence of large mucin pools and extravasated blood (Fig. 12C). In addition, adjacent reactive liver tissue frequently showed infiltration of inflammatory cells, fibrosis, or congestive sinusoids. Some tumor metastases showed abundant intratumoral collagen (Fig. 13A), while it was scarce in others (Fig. 13B-D). Different growth patterns were present at the CRC metastasis-liver parenchyma interface: *desmoplastic*, showing a wide band of collagenous tissue (Fig. 13B), separating the tumor and the liver parenchyma; *expansive*, lacking a collagen band, and metastases pushing away hepatocyte plates (Fig. 13C); and *infiltrating*, with a cell replacement pattern in which metastatic cells infiltrate the liver parenchyma without obvious alteration of the tissue structure (Fig. 13D).

**Figure 12.**
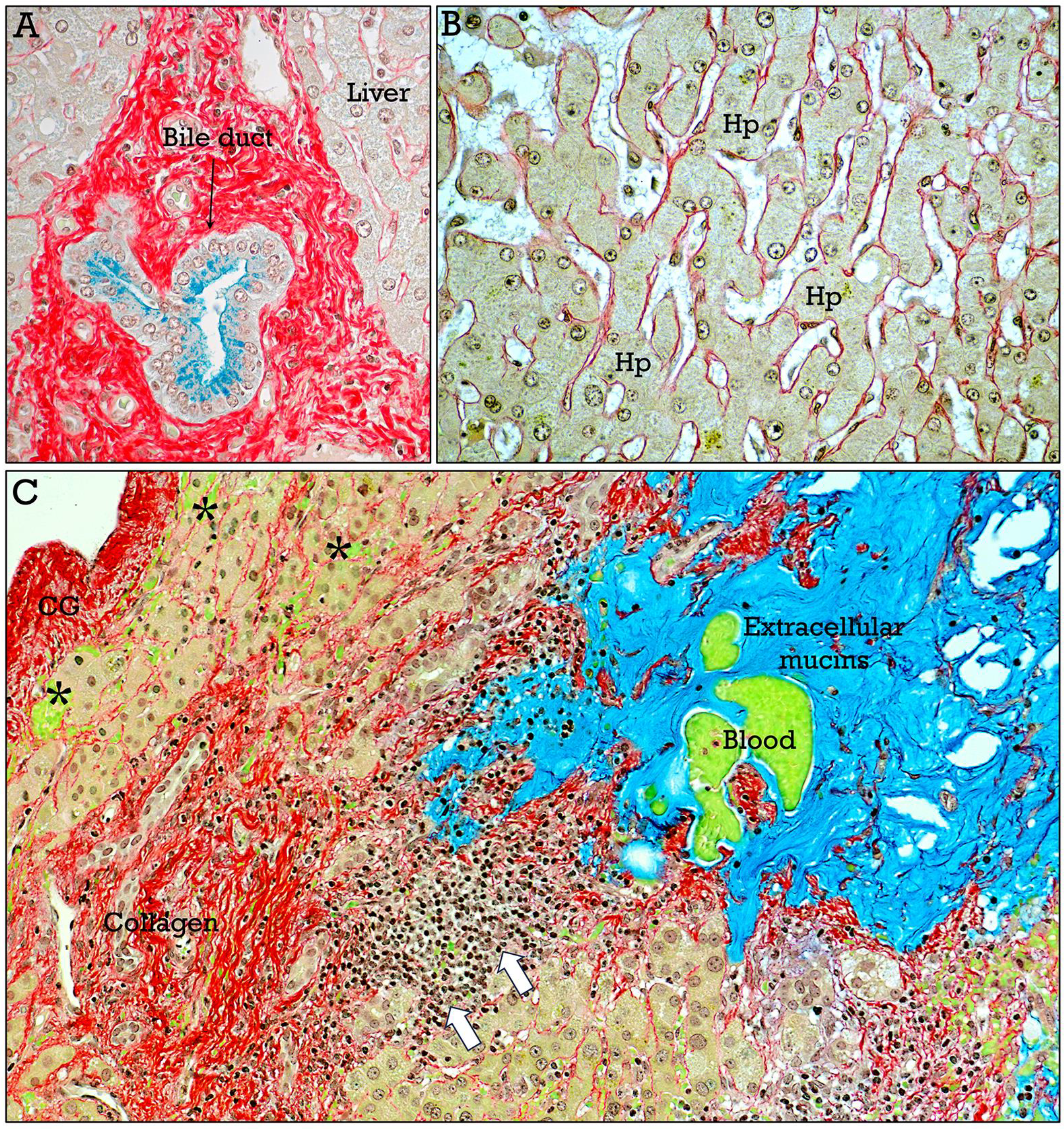
HemRGB-stained liver metastases (I). In normal liver, bile ducts (**A**) show apical epithelial mucin expression. Liver parenchyma (**B**) showing interconnecting hepatocyte plates (***Hp***) surrounding sinusoids. **C**, general view of a CRC metastasis showing areas of extracellular mucin, extravasated blood, inflammatory cell infiltration (***arrows***), increased collagen content, and vascular congestion (***asterisks***). ***GC*** capsule of Glisson.

**Figure 13.**
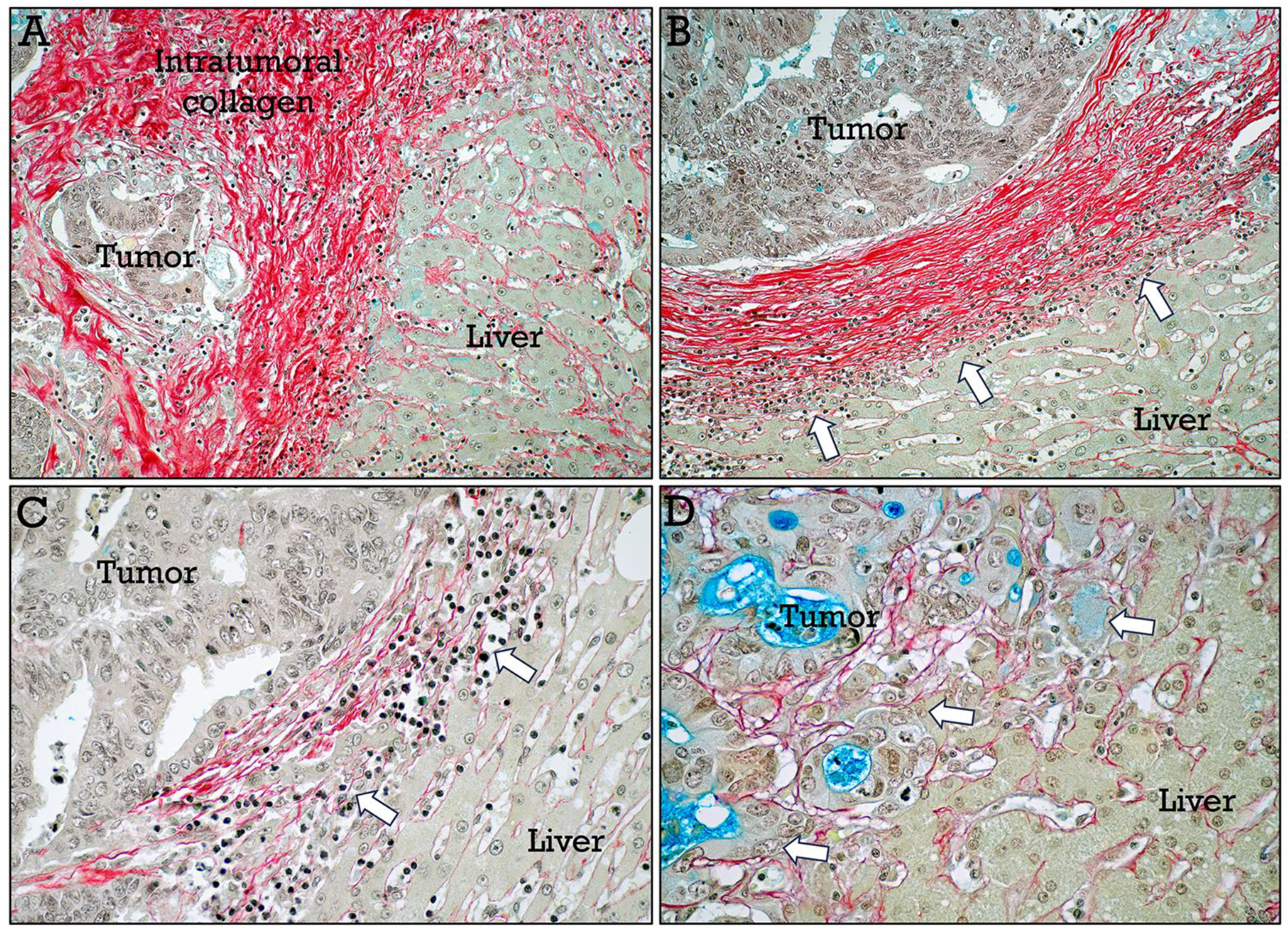
HemRGB-stained liver metastases (II). Interface between the CRC metastases and liver parenchyma. Large differences between cases were found with respect to the amount of intratumoral fibrosis (**A**), and different types of CRC metastasis-liver interfaces were observed: desmoplastic (**B**), expansive (**C**) and infiltrating (**D**), depending on the presence or absence of a band of collagenous tissue at the interface (***arrows***) or the direct infiltration of the liver

## Discussion

CRC is the third most common cancer in both sexes and is responsible for a large part of cancer-related mortality worldwide. Histopathological analysis is essential for the identification, classification, and grading of CRC cancer, providing crucial information for prognosis assessment^**7-9**^. Colorectal cancer evaluation relies on various histological features, including nuclear pleomorphism^**4**^, the degree of immune cell infiltration^**10**^, mucin secretion^**11**^, and tumor extracellular matrix characteristics^**12**,**13**^. The HemRGB trichrome addresses the limitations of RGB trichrome by enhancing nuclear staining and highlighting relevant histologic components, offering a comprehensive tool for CRC evaluation.

Nuclear pleomorphism, together with mitotic alterations, are hallmarks of cancer and have been reported to be related to CRC biology and clinical behavior^**3-5**^. Alterations of nuclear morphology and chromatin pattern can be easily identified in HemRGB-stained tissues. Nuclear staining is also useful to evaluate the presence of infiltrating leukocytes, primarily lymphocytes, that can be identified by their nuclear characteristics, and are likely related to the immune response against tumor cells^**10**^.

Mucin gene expression has been reported to be altered in CRC, and changes in mucin secretion have been proposed as a prognosis factor in CRC^**11**,**14**,**15**^. The data of this study agree with previous studies reporting a general decrease in mucin secretion except in some CRC subtypes with a mucinous component and showing abundant extracellular mucins^**11**^. The differential staining of some mucin-secreting cells in some tumors are likely due to the presence of both neutral (stained red) and acidic (stained blue) mucins, resembling the staining properties of mucins in metaplastic gastric mucosa expressing both neutral gastric and acidic intestinal mucins. This is in accordance with the reported dysregulation of mucin gene expression in CRC^**14**^.

The extracellular matrix is an essential component of the tumor microenvironment, and the amount of collagen and ground substance seems to be associated with cancer cell survival, proliferation, and invasive capacity^**12**,**13**^, acting either directly on tumor cells or indirectly through interaction with other parameters^**10**^. The HemRGB staining method is particularly reliable for the specific detection of collagen, that is the major structural component of the extracellular matrix, and has been reported to play important roles in tumorigenesis and cancer progression^**12**,**13**^. Accordingly, the abundance and distribution of collagen seem to be also related to the expansion and invasiveness of liver metastases^**16**,**17**^.

In addition to providing clear and specific staining of relevant histologic components of the CRC and the associated extracellular matrix, the HemRGB trichrome provides high-contrasted tissue interfaces, facilitating analytic procedures based on tissue segmentation, expanding the usefulness of the RGB trichrome as a tool for histopathological studies. Last but not least, and as stated by McGavin in a comprehensive review on the chromatic aspects of histological staining methods^**1**^, the nature of colors and the aesthetic characteristics of stained tissues have psychological effects on the viewer, likely affecting the time spent examining the sample and, therefore, the accuracy of tissue evaluation.

## Acknowledgements

The author is very grateful to Dr C Morales and Dr C Reymundo for advice, and to Dr E Tarradas for his technical support.

